# Wat1/mLst8, a TOR complex protein regulates mitochondrial integrity and calcium ion homeostasis in fission yeast *S. pombe*

**DOI:** 10.1101/2023.03.27.534337

**Authors:** Simmi Anjum, Swati Srivastava, Lalita Panigrahi, Uzair Ahmad Ansari, Arun Kumar Trivedi, Shakil Ahmed

## Abstract

The mTOR complexes play a fundamental role in mitochondrial biogenesis and cellular homeostasis. Wat1, an ortholog of mammalian Lst8 is an important component of TOR complex and is essential for the regulation of downstream signaling. Earlier we reported the role of Wat1 in oxidative stress response. Here, we show that the inactivation of *wat1* leads to respiratory defects and mitochondrial depolarization leading to decrease in ATP production. The confocal and electron microscopy in *wat1*Δ cells revealed the fragmented mitochondrial morphology implying its role in mitochondrial fission. Furthermore, we also showed its role in autophagy and the maintenance of calcium ion homeostasis. Additionally, *tor2-287* mutant cells also exhibit defects in mitochondrial integrity indicating the TORC1-dependent involvement of Wat1 in the maintenance of mitochondrial homeostasis. The interaction studies of Wat1 and Tor2 with Por1 and Mmm1 proteins revealed a cross-talk between mitochondria and endoplasmic reticulum through the Mitochondria-associated membranes (MAM) and endoplasmic reticulum-mitochondria encounter structure (ERMES) complex, involving TORC1. Taken together, this study demonstrates involvement of Wat1/mLst8 in harmonizing various mitochondrial functions, redox status, and Ca^2+^ homeostasis.

## Introduction

Eukaryotic organisms develop the ability to alter cellular metabolism in response to environmental changes, and mitochondria are the key organelle to mitigate these challenges (Sokolova 2018). Aerobic metabolism results in the continuous generation of reactive oxygen species (ROS) as a by-product of oxidative phosphorylation which are confined to the organelles like mitochondria and peroxisomes (Jayavelu et al., 2016). ROS homeostasis is maintained by an antioxidant system which is necessary for cell survival and signaling (Dickinson & Chang 2011). Excessive ROS production damages mitochondrial leading to a wide range of diseases like cancer, obesity, myopathies, and neurodegeneration (Balaban et al., 2005). A variety of stress conditions such as oxidative stress affects mitochondrial dynamics that trigger mitochondrial dysfunction (Xu et al., 2016; Guo et al., 2013). The dynamics of mitochondria as well as their morphology are regulated by many factors like mitochondrial fusion, fission events, and mitochondrial biogenesis (Meyer et al., 2017). However, damaged mitochondria produce an excessive ROS that creates a vicious cycle where more damaged mitochondria are created with concurrent ROS production.

TOR (Target of Rapamycin), is an evolutionarily conserved protein kinase that is the central controller of growth, metabolism and is involved in the maintenance of cellular homeostasis by sensing the nutrient and energy status of the cell (Loewith & Hall 2011). It plays a key role in coupling the mitochondria with cellular metabolism in a diverse range of organisms (Butow 2004; Liu & Butow 2006). Previous studies in mouse embryonic fibroblast cells have reported an accelerated rate of mitochondrial biogenesis and oxidative metabolism in response to genetic stimulation of mTORC1 (Cunningham et al., 2007). Additionally, an increase in ATP generation and mitochondrial respiration in skeletal muscle are associated with the synthesis of mTORC1-sensitive mitochondrial proteins (Morita et al., 2013). In muscle cells, the mTORC1 constitutive signaling activation increases mitochondrial respiration whereas its inhibition results in a metabolic switch from oxidative to glycolytic activity (Cunningham et al., 2007; Bentzinger et al., 2008; Morita et al., 2013). Interestingly, mTORC1 inhibition via rapamycin results in the switching of metabolism from glycolysis to activated amino acid catabolism in animals with diminishing mitochondrial function (Johnson et al., 2013). However, the deletion of mTORC2 signaling does not affect mitochondrial synthesis in muscle cells in comparison to mTORC1 but their oxidative metabolism is significantly decreased (Bentzinger et al., 2008; Risson et al., 2009). Furthermore, mTORC2-activated cells depend on aerobic glycolysis and mitochondrial oxidative phosphorylation for ATP production as revealed by a genome-wide shRNA screening (Colombi et al., 2011). The mitochondria-associated ER membrane (MAM) provides the site for the localization of mTORC2, and mTORC2-Akt-mediated calcium release at the MAM is important for mitochondrial physiology and ATP production (Betz et al., 2013; Singh et al., 2016).

Wat1, an ortholog of mammalian Lst8 is an important component of both TOR complexes TORC1 and TORC2 and regulates the downstream signaling (Ahamad et al., 2018, Panigrahi et al., 2023). Its role in genomic stability and microtubule integrity has also been reported (Verma et al., 2014; Ochotorena et al., 2001). Previously, we have shown that the inactivation of Wat1 in fission yeast cells results in ROS accumulation that might affect the mitochondrial integrity (Ahamad et al., 2016). In this study, we investigated the role of Wat1 in the maintenance of mitochondrial homeostasis. The disruption of *wat1* results in respiratory defects, mitochondrial depolarization, and reduction in ATP production. The confocal and electron microscopy in *wat1* knockout cells revealed fragmented mitochondrial morphology. Furthermore, its role in autophagy and calcium ion homeostasis has been elucidated involving ERMES and MAM complex.

## RESULTS

### Abrogation of *wat1* leads to respiratory defects and mitochondrial dysfunction

In a previous study, we showed that the deletion of *wat1* in *S. pombe* leads to ROS accumulation resulting in cell death (Ahamad et al., 2016). Growing evidences suggest that ROS generation mainly occurs in the mitochondria where it causes severe damage to the cell which is often associated with various pathologies (Murphy, 2009). To ascertain the role of Wat1 in maintaining mitochondrial integrity, the growth pattern of wild type and *wat1Δ* cells was examined in the media containing a non-fermentable carbon source (0.1% glucose and 3% glycerol). The *wat1Δ* and *wat1-17* mutant cells exhibit mild sensitivity in the media containing non-fermentable carbon source as compared to the wild type cells (Fig 1A). In comparison, these cells exhibit normal growth in complete rich (yeast extract) media containing 2% glucose (Fig 1A). The inefficiency of *wat1Δ* cells to use glycerol as a carbon source suggests that these mutant cells might have respiratory defects.

**Figure 1.**
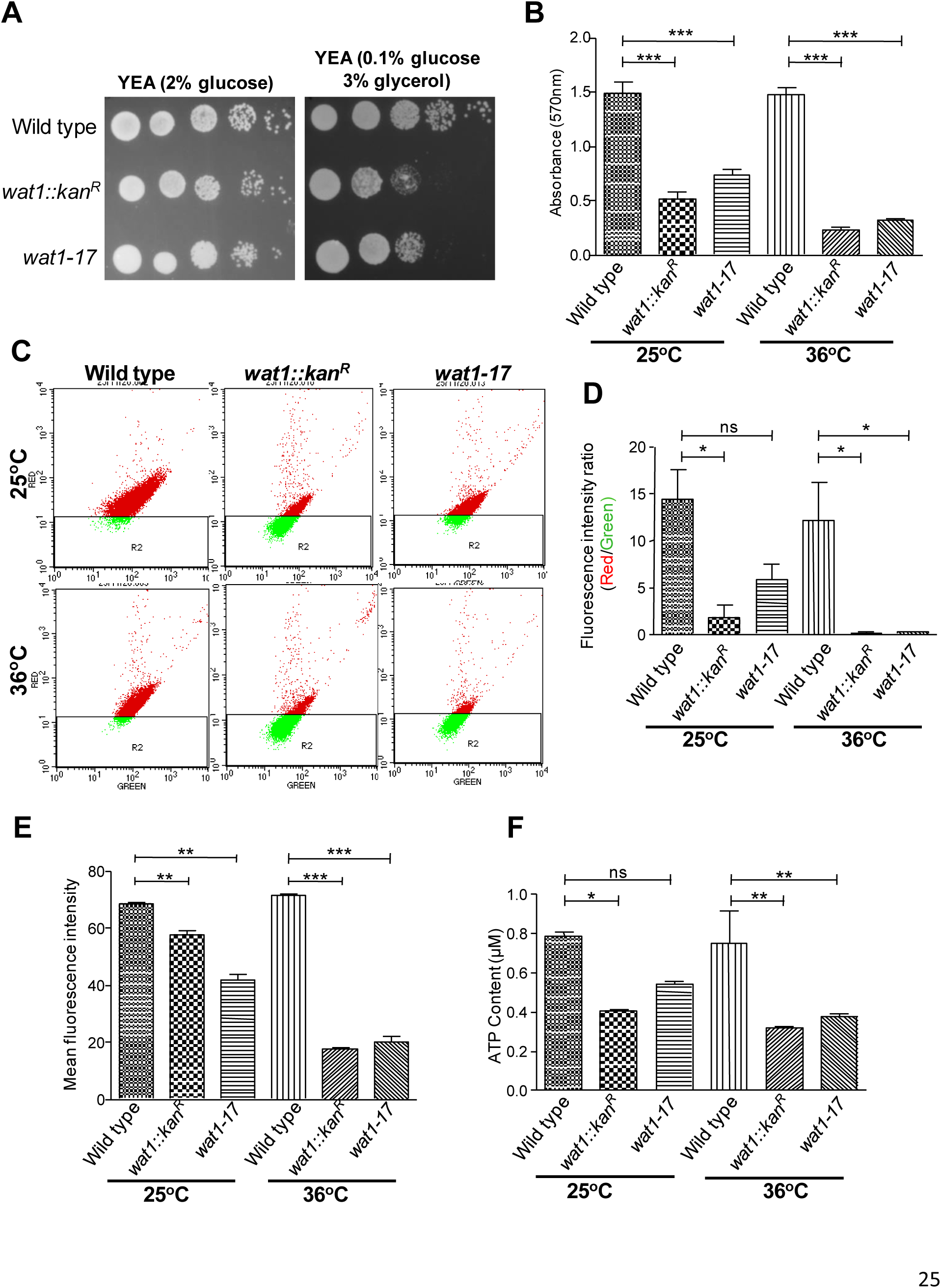
Abrogation of *wat1* affects the mitochondrial integrity. **(A)** Cells were grown till mid-log phase, serially diluted, and spotted on yeast extract medium containing 2% glucose or 0.1% glucose and 3% glycerol as a carbon source. Plates were incubated at 25°C for up to 4 days before taking photographs. **(B)** Wild type, *wat1*Δ, and *wat1-17* mutant cells were grown till mid-log phase and shifted at 36°C for 3h. Cells were processed with MTT as described in Materials and Methods. Absorbance was measured at 570nm and values were plotted. **(C)** Indicated strains were grown at 25°C till mid-log phase and shifted to 36°C for 3 h. The cells were collected, processed with JC-1 dye as described in Materials and Methods, and subjected to FACS analysis. **(D)** The ratio of cell population showing red and green fluorescence intensity was calculated and the values were plotted. **(E)** The cells were processed as above and incubated with rhodamine 123 dye for 30 min at 30°C, the mean fluorescence intensity was quantified by FACS and the values were represented as a bar graph. **(F)** The ATP content was measured as described in Materials and Methods. The experiment was performed in triplicate and the values were plotted as a bar graph. Asteriks show p values *p < 0.05, ** p < 0.01, ***p < 0.001.

The respiratory incompetency in *wat1Δ* cells along with the previously observed increase in the number of propidium iodide (PI) stained cells in a *wat1Δ* strain (Ahamad et al., 2016) led us to analyse the mitochondrial functionality using MTT assay. The cells having metabolically active mitochondria can cleave the tetrazolium ring of 1-(4,5-dimethylthiazol-2-yl)-3,5-diphenyl-tetrazolium bromide (MTT) using the enzyme succinate dehydrogenase and changes the colour of MTT from yellow to dark blue and hence has been used as an indicator of mitochondrial functionality and cell viability (Sanchez & Konigsberg 2006; Mosmann, 1983). MTT assay performed in the cells grown at permissive temperature, approximately three and two-fold reduction in the absorbance at 570nm was observed in *wat1Δ* and *wat1-17* mutant cells respectively as compared to wild type cells (Fig 1B), suggesting a defect in mitochondrial function. Interestingly, upon shifting the cells at non-permissive temperature, the absorbance was further reduced (4-6 fold) in *wat1Δ* and *wat1-17* mutant as compared to wild type cells grown under same conditions (Fig 1B). These results suggest that the abrogation of *wat1* may lead to a defect in the mitochondrial function which is further enhanced at non-permissive temperature.

### Dissipation of Mitochondrial Membrane Potential in the absence of *wat1*

The mitochondria are the major source of ROS and excess of it can damage the mitochondrial membrane, leading to the lowering of mitochondrial membrane potential (Simon et al., 2000). To elucidate the effect of excessive ROS production, the mitochondrial membrane potential was determined as an indicator of mitochondrial integrity. We sought JC-1 for evaluation of mitochondrial membrane potential in *wat1*Δ cells. In healthy mitochondria, it accumulates as an aggregate that fluoresces red; however, when the mitochondria are depolarized, it exists as a monomer and gives a green fluorescence. We observed a reduction in the JC-1 aggregates (red fluorescence) and an increase in the monomeric form of JC-1 (green fluorescence) in *wat1Δ* cells as compared to wild type cells (Fig 1C & D). In comparison, the *wat1-17* mutant cells did not exhibit any significant reduction in the JC-1 aggregates (red fluorescence) at permissive temperature (Fig 1C). At non-permissive temperature, there was a further reduction in the JC-1 aggregate (red fluorescence) leading to approximately 50 fold reduction in red/green fluorescence intensity ratio in *wat1Δ* and *wat1-17* mutant cells as compared to wild type cells (Fig 1C & D) suggesting the role of *wat1* in the maintenance of mitochondrial membrane potential. Additionally, the mitochondrial membrane potential was further evaluated by Rhodamine 123, a fluorescent dye which is quickly absorbed by healthy mitochondria resulting in an increase in green fluorescence. As shown in Fig 1E, at permissive temperature, a reduction in the fluorescence intensity was observed in *wat1Δ* and *wat1-17* mutant cells as compared to wild type cells. Upon shifting the cells at non-permissive temperature, the *wat1Δ* and *wat1-17* mutant cells exhibited further reduction (3-4 fold) in the fluorescence intensity while in wild type cells, it remained the same (Fig 1E). These results indicate mitochondrial dysfunction in *wat1Δ* and *wat1-17* mutant cells that lead to the lowering of mitochondrial membrane potential and hence reduction in average fluorescence intensity.

### The *wat1* disruption mitigates intracellular ATP level

The mitochondria play an important role in the production of ATP by generating proton motive force by an electrochemical proton gradient across the mitochondrial membrane (Noji et al., 1997). This electrochemical gradient which is essential for the synthesis of ATP is directed in such a way that it pushes the transport of cations toward the inner side and anions toward the outer side which eventually drives ATP production (Zorova et al., 2018). Mitochondrial dysfunction affects the mitochondrial permeability transition pore leading to leakage of solutes, dissipation of mitochondrial membrane potential, and decrease in ATP formation (Halestrap 2009). The mitochondrial depolarization in the absence of *wat1* led us to analyze the ATP content in these cells. As compared to wild type cells, we observed a twofold and 1.5-fold reduction in the total ATP content at permissive temperature in *wat1Δ* and *wat1-17* mutant cells respectively (Fig 1F). The total ATP content at non-permissive temperature further decreased by approximately 2.5 fold in *wat1Δ* and *wat1-17* mutant cells as compared to wild type cells (Fig 1F) suggesting that the *wat1* disruption affects the intracellular ATP content.

### Loss of *Wat1* induces a change in the morphology and mitochondrial dynamics

The mitochondria of almost all eukaryotic cells grow, divide, and fuse which governs the size, number, and shape of this dynamic organelle. Under normal conditions, they form inter-connected tubular elongated structures. The tubular network of mitochondria is essential for proper functioning which is achieved through the balance between fusion and fission events (Lackner 2014). We analyzed the mitochondrial dynamics by examining the morphological changes in the absence of *wat1* using GFP-tagged Cox4, a mitochondrial oxidase subunit. The confocal microscopic analysis revealed a tubular network of mitochondria in 80% of wild type cells at 25°C and 36°C (Fig 2A). In contrast, at permissive temperature only 25% of *wat1*Δ and 40% of *wat1-17* mutant cells displayed tubular mitochondrial morphology while 55% cells exhibited fragmented mitochondria as shown by small individual dots (Fig 2A & B). Interestingly, at non-permissive temperature, approximately 80% of *wat1*Δ and 60% of *wat1-17* mutant cells displayed fragmented mitochondria. These results indicate the role of Wat1 in maintaining mitochondrial dynamics.

**Figure 2.**
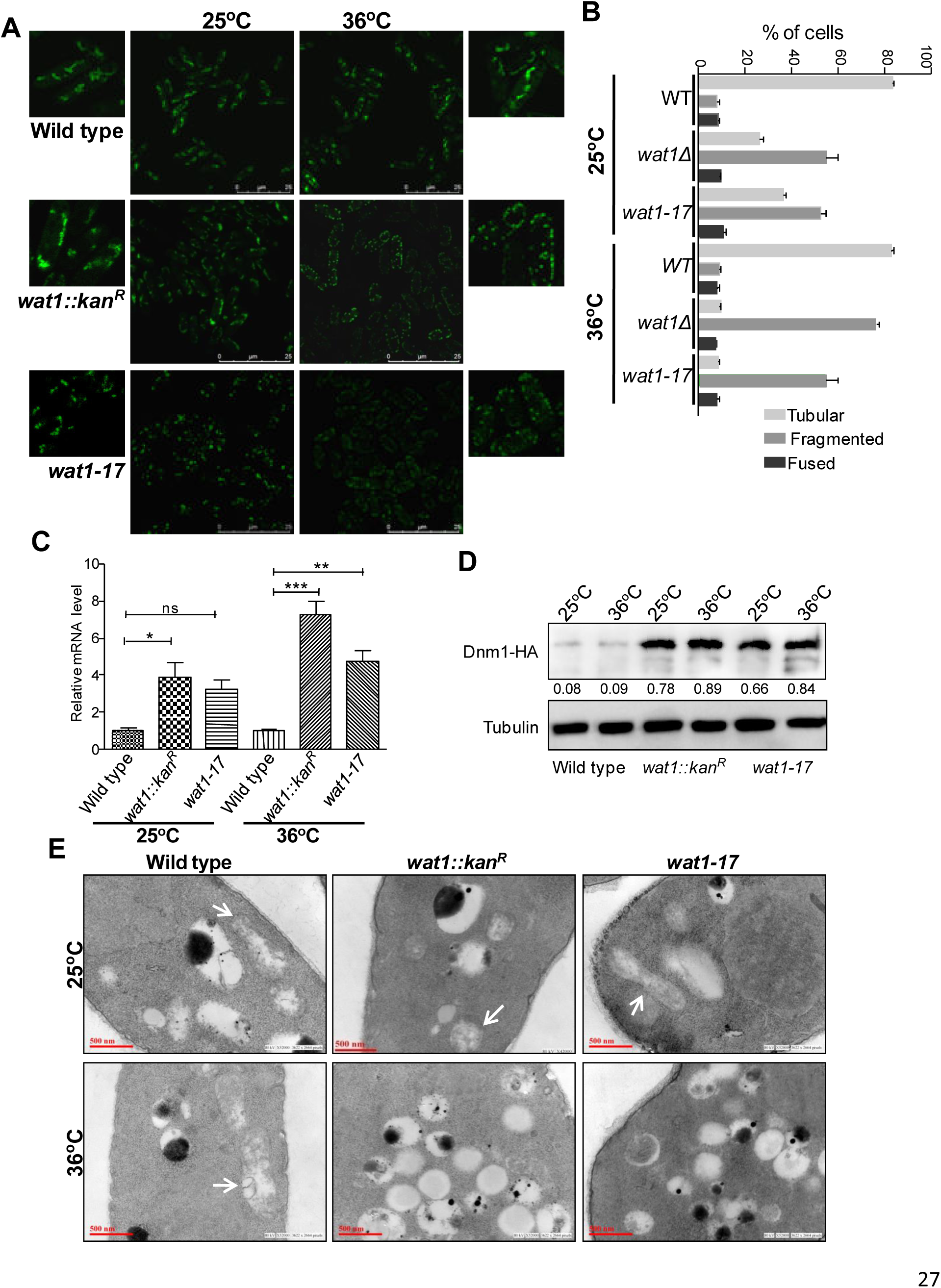
Loss of *wat1* results in mitochondrial fission and affects its morphology. **(A)** Indicated strains containing GFP tagged Cox4 were grown till mid-log phase and shifted to 36°C for 3hr. Cells were visualized using a confocal microscope **(B)** Approximately 100 cells for each strain were counted, the percentage of cells showing various mitochondrial morphology was calculated and plotted as a bar graph. **(C)** The cells were processed as described in Material and Methods, and RT-PCR analysis was performed using *dnm1* specific primers. The relative fold change for each strain was calculated and the average of values from three independent experiments was plotted. The asterisks indicate the p values * < 0.05, ** < 0.01, *** < 0.001. **(D)** The indicated strains were grown till mid-log phase and shifted to 36°C for 1hr. Protein lysate was prepared and western blot analysis was performed using an anti-HA antibody (Santacruz). The band intensity was measured using GelQuant Express Image analysis software and the ratio between Dnm1-HA and tubulin was indicated. **(E)** The cells were processed as described in Materials and Methods and Transmission Electron Microscopy (TEM) was performed to visualise the ultra-structural changes in *S. pombe* cells.

Furthermore, we also checked the expression of genes involved in mitochondrial fission (*dnm1*), and fusion (*fzo1*, *msp1*) in the absence of *wat1* by RT-PCR analysis. The cells grown at permissive temperature exhibited 4 and 3-fold increase in the expression of *dnm1* in *wat1Δ* and *wat1-17* mutant cells respectively as compared to wild type cells (Fig 2C). Upon shifting the cells at non-permissive temperature this increase in the expression of *dnm1* further enhanced to 7 and 5 fold in *wat1Δ* and *wat1-17* mutant cells respectively as compared to wild type cells (Fig 2C). The western blot analysis using Dnm1*-*HA tagged strain also revealed 7-11 fold increase in the expression of Dnm1 in *wat1Δ* and *wat1-17* mutant cells as compared to wild type cells at permissive as well as non-permissive temperature (Fig 2D). Consistently, the expression analysis of fusion genes, *fzo1* and *msp1* by RT-PCR analysis also revealed a decrease in the expression of these genes in *wat1Δ* and *wat1-17* mutant cells as compared to wild type cells (Supplementary Fig 1A & B). These results demonstrate that in the absence of Wat1, the imbalance between the expression of fusion and fission genes affects mitochondrial integrity.

Further to characterize the role of Wat1 in the structural maintenance of mitochondria, thin-section transmission electron microscopy (TEM) was performed in wild type, *wat1Δ,* and *wat1-17* mutant cells. We observed intact mitochondrial morphology in wild type cells at permissive and non-permissive temperatures (Fig 2E). In comparison, we observed only a minor loss of mitochondrial cristae in *wat1Δ* and *wat1-17* mutant cells at permissive temperature (Fig 2E). However, at non-permissive temperature the mitochondria mostly diminished in *wat1Δ* and *wat1-17* mutant cells with a dramatic increase in the number of vacuoles (Fig 2E) as have also been shown previously (Ahamad et al., 2018). These data indicate that the aberrant mitochondrial morphology may further lead to enhanced ROS formation which in turn disturbs the integrity of mitochondria.

### The absence of *wat1* leads to a decrease in mitochondrial DNA content

Previous studies have reported that the mitochondrial network dynamics govern the maintenance of mitochondrial DNA content (Delerue et al., 2016). The defect in mitochondria integrity was further analyzed by measuring the mitochondrial DNA content using qRT-PCR. The *cox1* and *cox2* are mtDNA-encoded genes and were used for the estimation of mitochondrial DNA content. In agreement with the loss of mitochondrial dynamics, we observed a decrease in mtDNA content in *wat1Δ* and *wat1-17* mutant cells at permissive temperature which was more profound at non-permissive temperature (Fig 3A & B). These results indicate that Wat1 plays an important role in the maintenance of mitochondrial volume.

**Figure 3.**
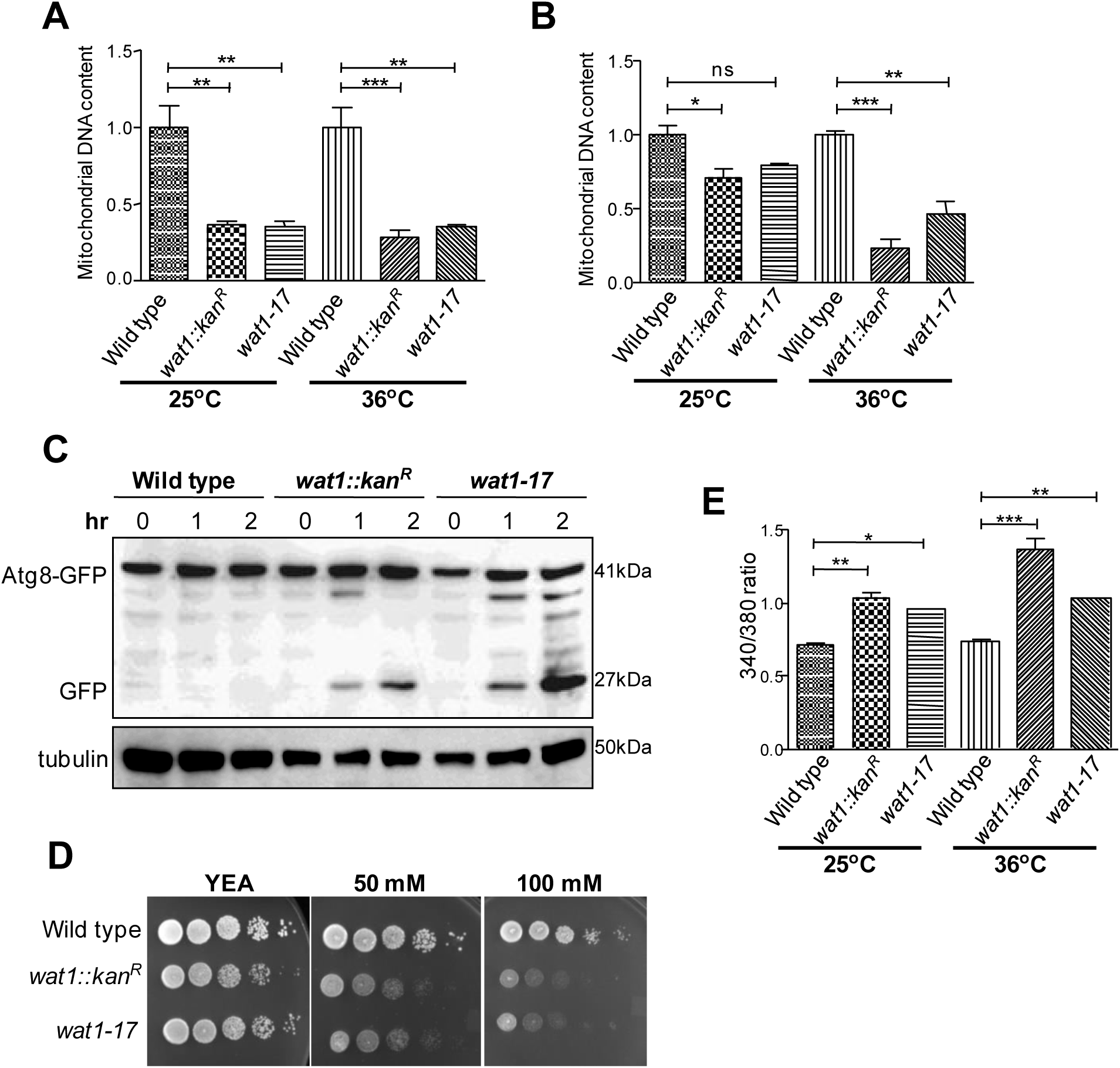
Inactivation of *wat1* results in the reduction of mitochondrial DNA content, induces autophagy and affects calcium ion homeostasis Indicated strains were grown till mid-log phase and shifted to 36°C for 3hr. DNA was extracted and subjected to qRT-PCR analysis using primers specific for mitochondria encoded genes *cox2* and *cox1*. The β-actin was used as a control. The level of mitochondrial *cox2* (A) and *cox1* (B) DNA relative to nuclear DNA from three independent experiments were calculated and represented as a bar graph. (C) The *S. pombe* cells containing Atg8-CFP were grown till mid-log phase and shifted to 36°C for indicated time points. Protein lysate was prepared and western blot analysis was performed using an anti-GFP antibody. (D) Indicated strains were grown till mid-log phase, serially diluted, and spotted on a plate containing 50 and 100mM CaCl_2_. Plates were incubated at 25°C for 4-5 days before taking photographs. (E) Indicated strains were grown till mid-log phase and shifted to 36°C for 3hr. Free cytosolic calcium level was calculated using Fura-2AM dye and a spectrofluorometer. The fluorescence intensity at 340 and 380nm was measured and the average ratio (F_340_/F_380_) from three independent experiments was represented as a bar graph. *p < 0.05,p**<0.01, ***p < 0.001.

### Loss of *wat1* induces autophagy

The mitochondrial permeability transition (MPT) leads to the changes in the outer mitochondrial membrane resulting mitochondrial degradation through macroautophagy (Gozuacik & Kimchi 2004). In general, the process of autophagy delivers Atg8-CFP from the autophagosome to the vacuoles where it is eliminated by proteolysis and replaced with free CFP which is stable and visible using an immunochemical assay (Yu et al., 2020). To investigate the role of Wat1 in autophagy, the processing of CFP-Atg8 fusion was investigated. We observed the continuous presence of CFP-Atg8 using anti-GFP antibody in all the samples at permissive and non-permissive temperatures (Fig 3C). Interestingly, the time-dependent accumulation of free CFP was observed only in *wat1Δ* and *wat1-17* mutant cells at non-permissive temperature which was absent in wild type cells (Fig 3C) suggesting that the absence of Wat1 might lead to autophagy.

### The disruption of Wat1 affects the calcium flux

It has been reported that under mild oxidative stress conditions, calcium ions releases from the ER accumulate in the cytosol (Ermak & Davies 2002). The functional mitochondria acts as a buffer organelle and helps in maintaining calcium ion homeostasis by transiently taking up calcium ions from cytosol thereby balancing the calcium ion concentration between mitochondria and cytosol (Panda et al., 2021). However, increased oxidative stress leads to cell membrane blebbing and cytoskeleton instability due to which calcium ion transfer is inhibited into the mitochondria (Stout et al., 1998). This results in an increase in cytosolic calcium ion concentration which along with oxidative stress can affect cellular growth (Orrenius et al., 1992). To envisage the role of Wat1 in maintaining Ca^2+^ homeostasis, we first performed a spotting assay to check the Ca^2+^ sensitivity of *wat1Δ* and *wat1-17* mutant strains. The wild type cells were able to form colonies on plates containing 50 and 100 mM concentrations of CaCl_2_. In comparison, we observed a dose-dependent CaCl_2_ sensitivity of *wat1Δ* and *wat1-17* mutant strains (Fig 3D) indicating the involvement of Wat1 in the maintenance of Ca^2+^ homeostasis. Furthermore, we also quantified intracellular calcium ion concentration using Fura2AM dye and monitored the fluorescence intensity ratio (F340/F380) by spectrofluorometer as described earlier (Betz et al., 2013). The results showed that in comparison to the wild type, there was a 1.5 and 1.3 fold increase in the intracellular calcium ions concentration in *wat1Δ* and *wat1-17* mutant cells respectively at permissive temperature (Fig 3E). However, at non-permissive temperature the Ca^2+^ concentration increases by 1.8 and 1.5 fold in *wat1Δ* and *wat1-17* mutant cells as compared to wild type respectively (Fig 3E). These findings raise the possibility that in the absence of *wat1*, calcium ion homeostasis is affected causing the release of mitochondrial content which along with the oxidative stress may lead to mitochondrial disruption.

### The *tor2-287* mutant cells exhibit defects in mitochondrial dynamics

The mammalian target of rapamycin complex 1 (mTORC1) has been implicated in regulating mitochondrial metabolism, translation of mitochondrial proteins, and management of mitochondrial turnover via balancing mitochondrial fusion and fission events (de la Cruz et al., 2019). Wat1, a mammalian homolog of mLst8 is a critical component of TOR complex and is required for the proper functioning of TORC1 and TORC2 (Panigarahi et al., 2023; Ahamad et al., 2018). These studies raise the possibility that the mitochondrial defects in the absence of Wat1 could be due to the non-functional TOR complex. To investigate this possibility, we checked the role of Tor2, a component of TORC1 in fission yeast, in the maintenance of mitochondrial integrity. The analysis of mitochondrial morphology using a GFP-tagged Cox4 revealed that the cells containing a temperature sensitive mutant allele of *tor2* (*tor2-287*) displayed fragmented mitochondrial morphology at permissive and non-permissive temperature (Supplementary Fig 2A) as was also observed in *wat1Δ* and *wat1-17* mutant cells. Furthermore, as compared to wild type cells, three and four-fold increase in the ROS generation was observed in *tor2-287* mutant cells at permissive and non-permissive temperature respectively (Supplementary Fig 2B) suggesting the mitochondrial dysfunctionality in the *tor2-287* mutant cells. The MTT assay revealed 1.7 and 3-fold reduction in the absorbance at 570nm in *tor2-287* mutant cells as compared to wild type cells at permissive and non-permissive temperature respectively (Supplementary Fig 2C). Interestingly, the *tor-287* mutant cells also exhibited a dose-dependent sensitivity on the plates containing CaCl_2_ (Supplementary Fig 2D). These results suggest that the mitochondrial defects in the absence of Wat1 might be due to the non-functional TORC1 pathway.

### Physical interaction of Wat1 with Por1 and Mmm1

Previous studies suggest that calcium ion signaling is responsible for dynamic crosstalk between ER and mitochondria through the ERMES complex to regulate oxidative stress (Murley & Nunnari 2016). ERMES complex consists of four components, Mmm1 (maintenance of mitochondrial morphology protein1), Mdm12 (mitochondrial distribution and morphology protein 12), and the OMM barrel proteins Mdm10 and Mdm34 (Kornmann et al., 2009). Growing evidence in mammalian cells suggests that the transfer of Ca^2+^ between ER and mitochondria is regulated by Ca^2+^ channels inositol tri-phosphate receptor (IP3R) of ER, VDAC1 (Por1 in *S. pombe*), and the cytosolic chaperone to maintain the calcium ion homeostasis (Drago et al., 2011; Poston et al., 2013). Therefore, to elucidate the mechanism of *wat1* in maintaining calcium ion homeostasis, we checked its plausible interaction with Por1/Vdac1 and ERMES complex protein Mmm1. Immunoprecipitation studies revealed a positive interaction of Wat1 with Por1 and Mmm1 proteins while Wat1-17 mutant protein was unable to interact with these proteins (Fig 4A & B). In agreement with the involvement of TORC1 in the maintenance of mitochondrial integrity and Ca^2+^ homeostasis, the Tor2 protein was also co-immunoprecipitated with Por1 (Fig 4C). Interestingly, in the absence of Wat1, the Tor2 was not able to interact with Por1 (Fig 4C) suggesting that the Wat1, an essential component of TORC1 is required for the interaction of Tor2 with Por1. Furthermore, we also observed the physical interaction of ERMES subunit Mmm1 with mitochondrial porin, Por1 (Fig 4D) as has also been shown in a large-scale affinity purification analysis in *S.cerevisae* (Kornmann et al., 2011; Murley et al., 2015). Interestingly, the interaction between Mmm1 and Por1 was completely lost in *wat1Δ* and *tor2-287* mutant cells (Fig 4D) suggesting that the TORC1 might be involved in facilitating this interaction.

**Figure 4.**
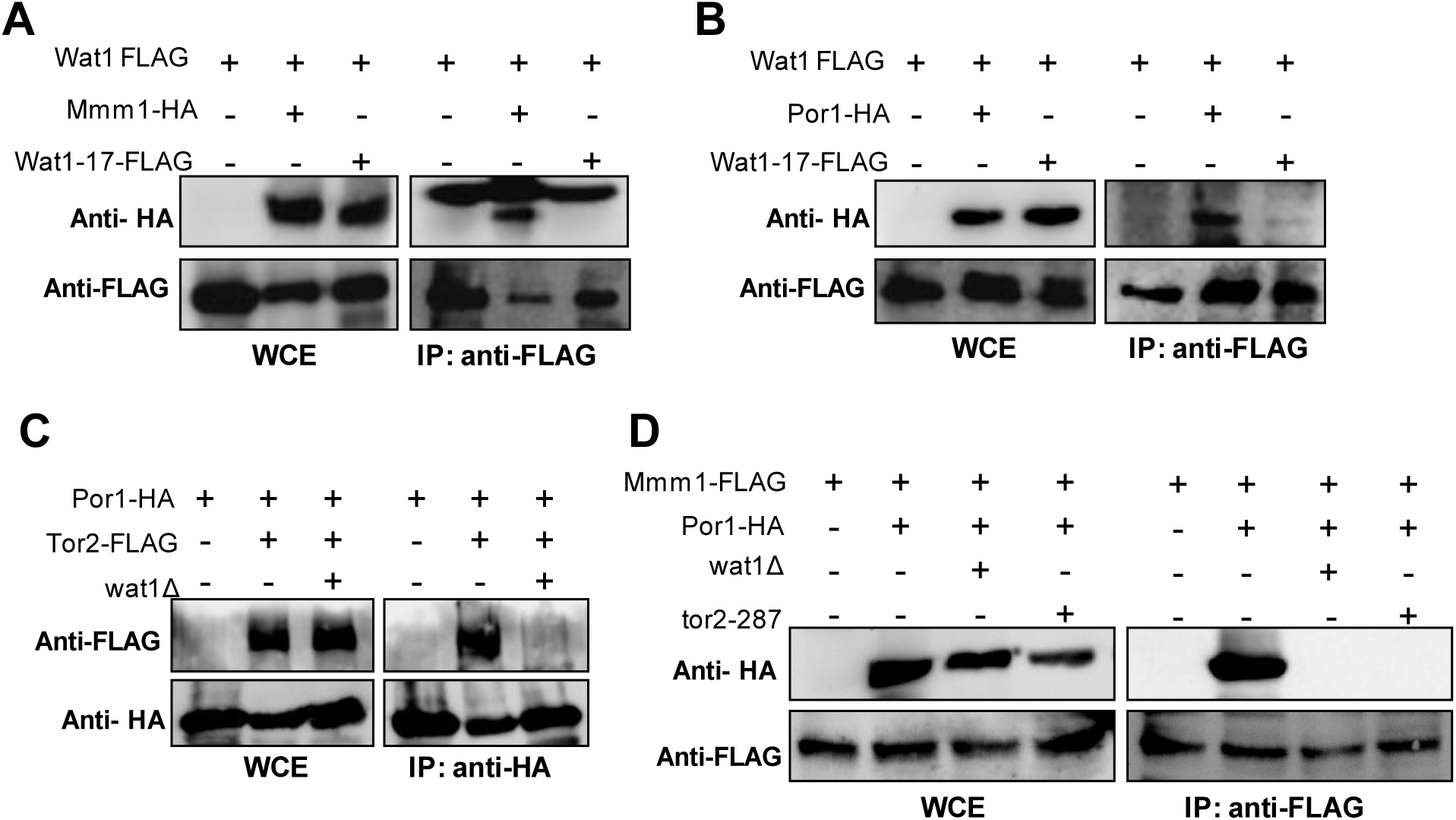
Interaction studies of Wat1 with Por1/Vdac1 and ERMES complex protein Mmm1. Protein lysate from indicated strains were prepared and co-immunoprecipitation between Wat1 and Mmm1 **(A)**, Wat1 and Por1 **(B)**, Por1 and Tor2 **(C)** and Mmm1 and Por1 **(D)** was performed as described in Materials and Methods. Samples were run on 8% SDS-PAGE, transferred on nitrocellulose membrane, and probed with anti-HA (F7) or anti-FLAG antibody.

### RNAi-mediated depletion of mLST8 (GβL) affects the mitochondrial integrity in HEK-293T cells

The mLst8 (also known as GβL) is a mammalian homolog of Wat1 protein. Based on our results in fission yeast, we investigated the role of mLst8 in the maintenance of mitochondrial integrity using the mammalian cell line (HEK-293T). Actively growing HEK-293T cells transfected with siRNA targeting mLst8 showed approximately 33% reduction in its expression compared to the cells transfected with control siRNA (Fig 5A). Flow cytometric analysis revealed 1.6 fold increase in ROS generation in the cells transfected with mLst8 siRNA as compared to cells transfected with control siRNA (Fig 5B). Furthermore, MTT assay performed in the cells transfected with mLst8 siRNA exhibited 1.5 fold reduction in the absorbance as compared to control siRNA (Fig 5C) indicating the malfunctioning of mitochondria under mLst8 knockdown condition.

**Figure 5.**
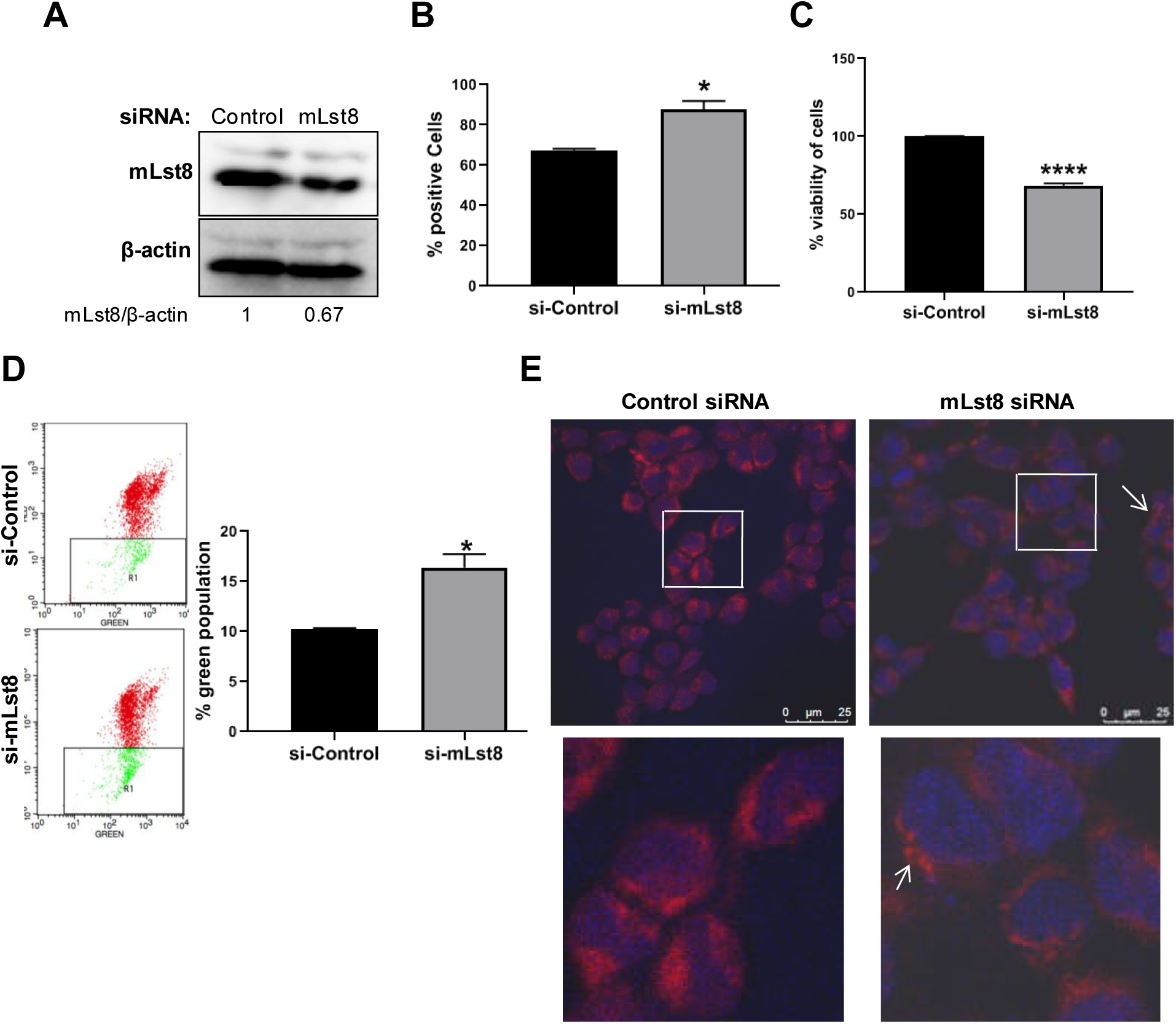
Knockdown of mLst8 alters the mitochondrial activity in HEK293T cells. Briefly, HEK-293T cells were seeded in 6 well plates. After 24 hrs, cells were transfected with either scrambled RNA (si-control) or siRNA for the mLst8 gene (si-mLst8). Cells were collected 48 hours post-transfection and assays were performed. **(A)** Cell lysates were collected and western blot was performed to confirm the knockdown of mLst8 gene. **(B)** Cells were washed with PBS followed by treatment with 10 μM DCFDA for 1 h. Cells were collected and FACS analysis was performed to estimate ROS generation. Histogram shows the ROS positive cells where n=2. **(C)** Cells were treated with 5mg/ml of MTT and incubated in dark for 4 hrs. Formazone precipitates were dissolved in DMSO and absorbance was measured at 570 nm. Histogram indicates the % viability of cells, where n=3. **(D)** Cells were washed with PBS followed by treatment with 10 μM of JC-1 for 1 hr. Cells were collected and FACS analysis was performed to estimate the red/green fluorescent intensity of cells. The left panel showed the representative image of red/green intensity. The right panel shows the quantitative analysis of % green cells, where n=3. **(E)** Cells were trypsinized and seeded on a coverslips followed by 12h of incubation. 100 nM concentration of mitotracker solution was added and samples were incubated at 37°C for 5 minutes. Cells were fixed in 4% PFA in PBS followed by immunofluorescence. Images were taken as 100X, where the scale bar is 25µm. All data are represented as mean ± SD of triplicates, where *, **, and **** indicate *P*<0.05, *P*<0.01, and *P*<0.0001 respectively.

Mitochondrial membrane potential analysis using JC-1 dye showed an increase in the monomeric form of JC-1 (green fluorescence) in mLst8 siRNA transfected cells as compared to control siRNA (Fig 5D) suggesting the lowering of mitochondrial membrane potential. The confocal microscopic analysis, using MitoTracker Red CMXros revealed abnormal punctate morphology in mLst8 knockdown cells as compared to continuous mitochondrial morphology in the control cells (Fig 5E) indicating the alteration in mitochondrial morphology. Unlike *S. pombe* knockout cells, we could not observe clear fragmented mitochondrial morphology in mammalian cells after siRNA knockdown. This could be due to the fact that after siRNA transfection, there was only a 33% loss of mLst8 expression in mammalian cells. Taken together, these results suggest the role of mLst8 in the maintenance of mitochondrial integrity in mammalian HEK-293T cells.

## Discussion

### The Wat1/mLst8 is required for maintaining the mitochondrial dynamics

The present study has attributed the effect of *wat1* disruption on mitochondrial homeostasis. Several lines of evidence suggest an intricate interplay between TOR signalling and mitochondrial integrity (Schieke & Finkel 2006; de la Cruz et al., 2019) but specifically the role of Wat1/mLst8, a component of TOR complex has not been explored. Our previous study provided evidence that *wat1* deficiency leads to oxidative stress and cell death due to excessive production of ROS in *S. pombe* (Ahamad et al., 2016). Additionally, it has been suggested that mitochondria serves as a primary target as well as a key generator of ROS (Wang et al., 2017). It is the main organelle responsible for mitigating the oxidative stress generated due to the accumulation of ROS. The inefficiency of *wat1*Δ and *wat1-17* mutant cells to grow on non-fermentable carbon source could be due to the malfunctioning of respiratory chain complexes of mitochondria that would further enhance ROS production and affect the mitochondrial integrity. Consistent with the temperature sensitive phenotype of *wat1Δ* and *wat1-17* mutant, these defects are more severe at non-permissive temperature as compared to permissive temperature. Additionally, an increase in the monomeric form of JC-1 dye in *wat1Δ* and mutant cells indicate the lowering of mitochondrial membrane potential and consequently a decrease in ATP production. It has been reported that the electrochemical proton gradient across the mitochondrial membrane regulates the movement of cations and anions which eventually drives ATP production (Zorova et al., 2018). Under normal conditions, the cells maintain the levels of intracellular ATP and mitochondrial membrane potential, and prolonged perturbation of these factors affects cell viability (Zorov et al., 2014; Izyumov et al., 2004). The severe loss of viability upon inactivation of Wat1 in *S. pombe* cells along with changes in mitochondrial membrane potential and reduction in ATP production suggests a role of Wat1 in the maintenance of mitochondrial integrity. In agreement, the electron microscopic analysis revealed mostly diminished mitochondria in *wat1Δ* and *wat1-17* mutant cells at non-permissive temperature with a dramatic increase in the number of vacuoles as have also been observed previously (Ahamad et al., 2018).

The confocal microscopic analysis using Cox4-GFP revealed the presence of small individual dots-like structures as compared to tubular mitochondrial morphology in *wat1Δ* cells suggesting the disruption of mitochondrial dynamics leading to mitochondrial fragmentation. The mitochondrial homeostasis network is maintained by a highly regulated event of fission and fusion, mediated by dynamin-related protein Dnm1 (Drp1 in mammals) and Fzo1 respectively (Bleazard et al., 1999; Smirnova et al., 2001; Yang et al., 2021). In agreement with these results, we also observed an increase in the transcript level of *dnm1* after the inactivation of Wat1 while the expression of fusion genes *fzo1* and *msp1* was reduced. Our results suggest that in the absence of *wat1*, the increase in ROS level and reduction in membrane potential could lead to mitochondrial fragmentation as well as a reduction in total mitochondrial DNA content.

### The role of fission yeast TORC1 in regulating the mitochondrial integrity and autophagy

The mammalian, as well as yeast TOR kinase protein, is present in two multi-protein complexes, TORC1 and TORC2. The mLst8, a mammalian ortholog of Wat1 is an important component of both complexes and can interact with the kinase domain of mTOR independently of the other factors (Yang et al., 2013). The mTORC1 also regulates the bioenergetics processes which are controlled by mitochondria (Dowling et al., 2010) and its role in regulating mitochondrial biogenesis, mitophagy, and apoptosis has been well reported (de la Cruz et al., 2019). Since the *tor2-287* mutant (a component of TORC1) also exhibits defects in mitochondrial integrity, we postulate that the Wat1 may play a role in the maintenance of mitochondrial homeostasis through the TORC1 complex. Consistently, the disruption of mLst8 by siRNA in mammalian cells also leads to defects in mitochondrial activity (Fig 5). Earlier studies have also demonstrated the disruption of TORC1 by rapamycin or RNAi affects the mitochondrial metabolism (Schieke et al., 2006). The mitochondrial degradation is regulated by mTORC1 by suppressing autophagy, consequently, the mTORC1 inhibition activates autophagy (Morita et al., 2013). In general, the process of autophagy delivers Atg8 from the autophagosomes to the vacuoles where it is eliminated by proteolysis. Consistently, we observed time-dependent proteolysis of Atg8-CFP in *wat1Δ* and *wat1-17* mutant cells at non-permissive temperature (Fig 3C).

### The regulation of calcium flux by Wat1 is mediated by MAM and ERMES complex involving TORC1 complex

The mitochondria and endoplasmic reticulum play an essential role in maintaining the Ca^2+^ homeostasis that involves calcium influx, efflux, and storage (Tang et al., 2015). An increase in cytosolic calcium ion concentration along with oxidative stress affects cellular growth (Orrenius et al., 1992). Increased sensitivity to calcium ions and increased cytosolic calcium ion concentration in the absence of *wat1* may be due to the malfunctioning of protein complexes that regulate calcium ion homeostasis under oxidative stress.

The mitochondrial permeability transition pore (MPTP) present on the inner mitochondrial membrane is involved in mitochondrial calcium homeostasis (Bernardi, 1999). The cytosolic decrease in Ca^2+^ results in the movement of mitochondria towards ER forming ER-mitochondrial contacts through mitochondrial associated membranes (MAMs) that leads to mitochondrial Ca^2+^ uptake (Csordas et al., 1999; Rizzuto et al., 1998). Numerous channels like inositol 1,4,5-trisphosphate (IP_3_) receptor (IP_3_R) Ca^2+^ release channel, mitochondrial outer membrane Ca^2+^ channel VDAC1 and the cytosolic chaperone Grp75 coordinate to transfer calcium ions between mitochondria and ER (Duchen 1999, Drago et al., 2011). In yeast, the ERMES complex containing four core proteins Mmm1, Mdm12, Mdm10, and Mdm34 form a molecular bridge between the ER and mitochondria (Kundu & Pasrija 2020). The Mmm1 containing a trans-membrane domain is an ER anchoring protein; Mdm12 is a cytoplasmic junction protein while Mdm10 and Mdm34 are outer mitochondrial membrane proteins. The ERMES complex acts as a mediator of lipid exchange between the two organelles (Lang et al., 2015). This study provides evidence of Tor2 and Wat1 dependent interaction of Mmm1 with Por1, a mammalian ortholog of VDAC1 that suggest a cross talk between ERMES complex and Ca^2+^ channel in fission yeast. The physical interaction of ERMES subunit Mmm1 with mitochondrial porin, Por1 has also been reported previously (Kornmann et al., 2011; Murley et al., 2015). All together, we summarize that under normal conditions the Ca^2+^ homeostasis is maintained by ERMES and MAM complex involving the TORC1 components Tor2 and Wat1. In the absence of Wat1 (or in tor-287 mutant), the integrity of EREMS and MAM complex is affected which leads to an increase in the level of Ca^2+^ in the cytosol. The ROS accumulation in the absence of Wat1 (Ahamad et al., 2016) along with increased Ca^2+^ affects the mitochondrial integrity and cellular growth (Fig 6). Although, the involvement of other components of TORC1 in this process cannot be ruled out. Additionally, the effect of RNAi-mediated depletion of mLST8 on mitochondrial integrity in HEK-293T cell line suggests the functional conservation of pathway in yeast and mammalian system. Overall, this study demonstrates the involvement of Wat1/mLst8 in harmonizing various mitochondrial functions including mitochondrial metabolism, biogenesis, and redox status which can be utilized for the development of anti-cancer therapeutics.

**Figure 6.**
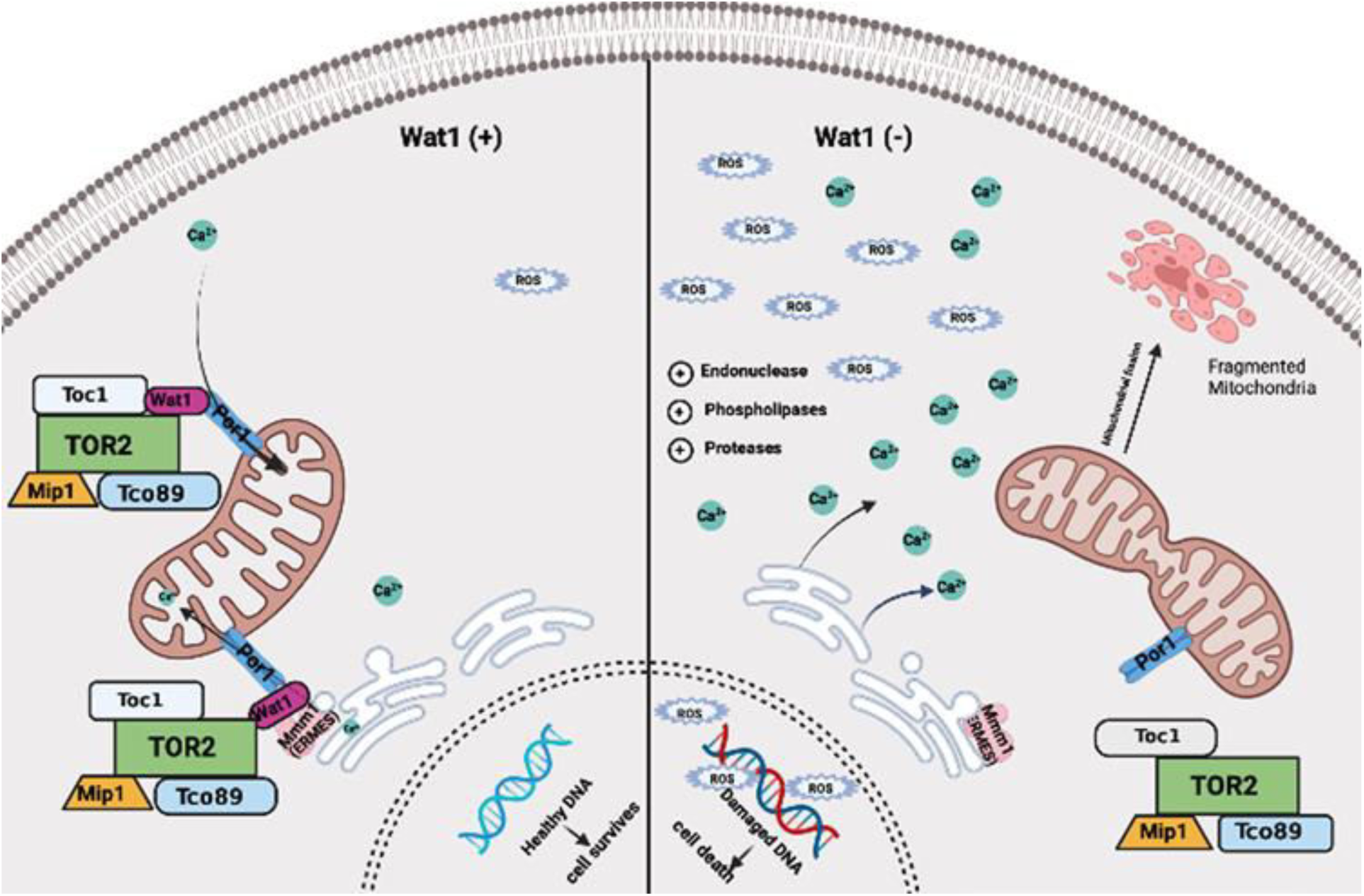
Mechanism underlying the contribution of Wat1 in the maintenance of mitochondrial integrity and Ca^2+^ homeostasis. Overproduction of ROS in the absence of *wat1* affects mitochondrial integrity. Under normal conditions, Ca^2+^ fluxes are regulated across the plasma membrane and between intracellular compartments. Excessive ROS generation, damage the proteins which regulate various Ca^2+^-regulating proteins resulting in an increase in the levels of Ca^2+^ in the cytosol. The interaction of TORC1 with Por1 and Mmm1 is responsible for maintaining the calcium ion transfer between ER and mitochondria. In the absence of Wat1, this loss of interaction leads to excessive accumulation of calcium ion in the cytosol and affect Ca^2+^ homeostasis.

## Materials and methods

### Strains preparation and media

The *S. pombe* strains used in this study are presented in Supplementary Table S1. The strains were prepared using standard genetic methods. For vegetative growth of the cells, the complete rich media containing 0.5% yeast extract, 2% glucose, and 100µg/ml adenine was used. For temperature shift experiments, cells were allowed to grow till the mid-log phase at 25°C and then shifted to 36°C for the desired time points For the spotting assays, approximately 10^7^ cells were serially diluted and spotted on the rich media plate. For respiratory defect analysis, the wild type and mutant cells were spotted on plates containing non-fermentable carbon source (0.1% glucose and 3% glycerol).

### Mammalian cell culture and siRNA transfection

HEK-293T cells were procured from ATCC and grown in DMEM-High glucose medium supplemented with 10% fetal bovine serum (FBS), 100 units/ml penicillin, 100 μg/ml streptomycin and 2 mM glutamine (Thermo Fisher Scientific, Waltham, MA). Scrambled RNA (si-Control) and siRNA for the mLst8 gene (si-mLst8) were purchased from Santa Cruz Biotechnology. Briefly, HEK-293T cells (1x10^6^) were seeded in a 6-well plate and the next day, cells were transfected with 50 nM of siRNA using oligofectamine reagent (Thermo Fisher Scientific) as per manufacturer’s instructions. After 48 hrs of transfection, cells were harvested and processed for various assays.

### Reactive Oxygen Species (ROS) generation assay

ROS generation in yeast cells was estimated using DCFDA (2’,7’–dichlorofluorescein diacetate) dye, where, DCFDA was oxidized by cellular ROS into 2’,7’–dichlorofluorescein. DCF is highly fluorescent and can be detected by fluorescence spectroscopy with excitation/emission at 485/535nm. For ROS estimation in mammalian cells, si-RNA transfected HEK-293T cells were treated with a 10 µM concentration of DCFDA solution and incubated for 1h in dark. Cells were washed and collected in PBS for FACS analysis using Flow Cytometer (BD Biosciences).

### MTT assay to measure Mitochondrial functionality

MTT assay to determine the mitochondrial functionality in *S.pombe* cells was performed as previously described (Sánchez & Königsberg 2006). Briefly, approximately 0.5x10^6^ cells were taken in 500ml of PBS, 50µ l of Methylthiazolyldiphenyl-tetrazolium bromide (MTT) was added and the mixture was incubated at 37°C for 3h in dark. The reduction of MTT by mitochondrial succinate dehydrogenase was determined using a visible spectrophotometer at 570 nm.

In mammalian cells, siRNA-transfected cells were seeded in a 96-well plate. After 24h, 5mg/ml of MTT solution was added to the cells followed by 4h incubation in dark. The formazan precipitate was dissolved in DMSO and absorbance was taken at 570 nm.

### Mitochondrial membrane potential measurement

Mitochondrial membrane potential (ψ_m_) was measured by using fluorescent probes JC-1 and Rhodamine 123 (Sigma Aldrich). For yeast and mammalian cells, approximately 1x10^7^ cells were washed with PBS, and 5µM or 10 µ M of JC-1 dye was added respectively, followed by incubation at 37°C for 30 min. After washing with PBS, the cells were subjected to FACS analysis. For rhodamine 123 staining, after washing the cells, 25µ M of rhodamine dye was added and incubated at 30°C for 30 min. Samples were processed as mentioned above and fluorescent intensity was quantified by FACS analysis.

### Measurement of mitochondrial DNA content

For mitochondrial DNA content measurement, *S. pombe* cells were grown till mid-log phase and shifted at 36°C for 3h. The DNA was isolated and approximately 25ng DNA was taken for real-time q-PCR experiment using the Biorad CFX C1000 thermal cycler real-time system and mitochondrial gene-specific primers (Supplementary Table S2). The β-actin gene was used as a nuclear DNA control. Relative nuclear to mitochondrial DNA level was calculated and the graph was plotted.

### Assessment of ATP concentration

*S. pombe* cells were grown till mid-log phase and shifted at 36°C for 3h. Protein lysate was prepared using FastPrep-24 (MP Biomedicals). 50 µ g of lysate was used for the assessment of ATP concentration using an ATP Determination Kit (ATP lite assay, PerkinElmer) as described earlier (Miyoshi et al., 2006). Briefly, 50µ g of cell lysate was mixed with a reaction buffer containing 1 mM dithiothreitol (DTT), 0.5 mM luciferin, and 12.5 μg/ml of luciferase. After gentle mixing of the solutions, the intensity readings were measured using Microplate Reader (Tecan, M200). The ATP concentration was calculated using an ATP standard curve.

### Mitochondrial Morphology Analysis

*S.pombe* strain containing the GFP tag Cox4 (Chacko et al., 2019) was used as a mitochondrial marker to assess the mitochondrial morphology. The GFP-tagged *cox4* gene was incorporated in desired strains by standard genetic crosses. The Cox4-GFP was visualized as a mitochondrial marker by confocal microscopy. The percentage of cells showing various mitochondrial morphotypes was calculated.

For mammalian cells, the mLst8 silenced cells were seeded on a cover slip. After 12h of incubation, cells were washed with PBS and treated with 100 nM Mitotracker solution for 5 minutes. Cells were fixed with 4% paraformaldehyde in PBS followed by immunofluorescence imaging. Images were taken as 100X, where the scale bar is 25 µ m.

For ultrastructural analysis, Transmission electron microscopy (TEM) was performed as described previously (Unger et al., 2017). Briefly, cells were centrifuged at 6000 rpm for 5 min, fixed with 4% paraformaldehyde at room temperature for 30min. After fixation, cells were washed with sorbitol buffer (1M sorbitol, 10mM MgCl_2_, pH7.8), and 6% osmium tetroxide was added followed by spheroplasting using zymolyase (1µg/ml) at 37°C for 30 min. Cells were centrifuged and the pellet was washed with sorbitol buffer followed by treatment with 1% uranyl acetate at room temperature for 4h. Dehydration was performed with different grades of ethanol (25, 50, 75, 95, and 100 %) for 5 min in each solution. Epoxy resin was used for embedding. Thin sections were cut and mounted on grids for observation in a transmission electron microscope (FEI Tecnai G-2 spirit).

### RNA extraction and cDNA synthesis

Cells were grown till mid-log phase and shifted at 36°C for 3h. Cells were collected and a hot phenol method was used to extract the RNA from yeast cells as described earlier (Sonkar et al., 2018). 1µg of RNA was reverse transcribed using a cDNA synthesis kit (Thermo Scientific) followed by qRT-PCR with SYBR green master mix kit (Takara). The reactions were performed in triplicate using the Agilent Mx3000P qPCR system and analysis was performed essentially as described earlier (Panigrahi et al., 2023).

### Measurement of calcium ion concentration

Cells were grown till mid-log phase and shifted at 36°C for 3h. Intracellular calcium ions were measured using calcium indicator dye Fura-2 AM (Betz et al., 2013). Cells were mixed with 10 mM Fura-2 AM in dark for 30 min. After washing, the cells were suspended in HEPES buffer and excitation spectra at 380 nm (calcium free) and 340 nm (calcium complex) were monitored with fixed emission at 510 nm using a spectrofluorimeter. The relative amount of free intracellular calcium was expressed as a change in the ratio of intensities following excitation at 340 nm to 380 nm.

### Preparation of whole cell lysate and Western blot analysis

The *S. pombe* cells were grown till mid-log phase at 25°C. Cells were collected by centrifugation and lysed using glass beads and a FastPrep-24 (MP Biomedicals). Lysate was prepared in Phosphate Buffered Saline (PBS) containing 200mM PMSF, 0.5% Triton X-100, and 1X Protease inhibitor cocktail (Sigma Aldrich). The mammalian cell lysate was prepared in RIPA lysis buffer containing 50 mM HEPES, pH 7.5, 10% glycerol, 150 mM NaCl, 1% Triton X-100, 1 mM EDTA, 1 mM EGTA, 10 mM NaF, and 30 mM β-glycerol phosphate. Samples were centrifuged at 12000 rpm in a microcentrifuge for 10 min at 4°C. The supernatant was collected and protein estimation was performed using the Bradford protein assay. For western blot analysis, 300 µ g of total cell lysate was run on 8% SDS-PAGE, transferred to nitrocellulose membrane (Amersham) using 10% methanol in transfer buffer with a constant current at 250mA for 4-5 h at 4°C. Immunoblotting was performed using anti-HA (1:1000 SantaCruz), anti-Flag (1:1000; Sigma Aldrich), or anti-mLst8 (1:1000, Santa Cruz) antibodies. A peroxidase-coupled secondary antibody and the enhanced chemiluminescence detection system (Millipore) were used to detect the immune complexes using Image Quabt LAS 4000 biomolecular imager (GE Healthcare).

### Co-immunoprecipitation

Soluble protein lysates from *S. pombe* cells were prepared in lysis buffer (PBS) containing 50 mM NaF, 1 mM PMSF, and 1X protease inhibitor. 600μg of protein was used for immunoprecipitation as described earlier (Ranjan et al., 2014).

### Statistical analysis

The data were analyzed using Graph pad prism 5.0 software (Lo Lalla, CA) and represented as ± the standard error of the mean (SEM) The data was obtained by performing at least three independent experiments. One-way ANOVA test followed by Turkey’s multiple comparison tests and Dunnett tests were used for comparing multiple experimental groups. p value denotes the significance.*p<0.05,**p<0.01,***p<0.001, ns –non significant.

## Acknowledgement

We thank the members of our lab for helpful discussions and technical support and; the yeast resource center NBRP, Japan for providing the strains as mentioned in the strain list. We thank Prof. Phong Tran and Vaishnavi Ananthanarayanan for Cox-GFP and Prof Li Lin for the Atg8-CFP strain. Our special thanks to the Sophisticated Analytical Instrument Facility (SAIF) of CDRI for FACS analysis. Electron Microscopic studies were performed in collaboration with the Indian Institute of Toxicological Research, Lucknow. S Anjum and UA Ansari acknowledge the Academy of Scientific and Innovative Research (AcSIR), Ghaziabad, 201002, India.

## Author Contributions

S Anjum designed and performed the experiments, and wrote the manuscript draft. S Srivastava performed mammalian cell line studies. L Panigrahi constructed the yeast strains and helped in real-time PCR experiments; UZ Ansari performed EM studies, S. Ahmed and AK Trivedi conceived the idea, supervised whole study, and wrote final manuscript.

## Funding

This work was supported by the grants from the Department of Biotechnology and Council of Scientific and Industrial Research, New Delhi, India. S. Anjum and L. Panigrahi acknowledge University Grant Commission (UGC) for providing a research fellowship.

## Availability of data and material

All the data generated during this study are included with in the article and supplementary materials. Raw images and raw data are available from the corresponding author on reasonable request.

## Ethics Approval and consent to participate

Not applicable

## Conflict of Interest

The authors declare that they have no conflicts of interest with the contents of this article.

## Notes

### Competing Interest Statement

The authors have declared no competing interest.

